# Insect herbivory and avian insectivory in novel native oak forests: divergent effects of stand size and connectivity

**DOI:** 10.1101/558114

**Authors:** Elena Valdés-Correcher, Inge van Halder, Luc Barbaro, Bastien Castagneyrol, Arndt Hampe

**Affiliations:** BIOGECO, INRA, Univ. Bordeaux, 33610 Cestas, France; Dynafor, INRA-INPT, Univ. Toulouse, Auzeville, France; CESCO, Muséum national d’Histoire naturelle, CNRS, Sorbonne-Univ., Paris, France

**Keywords:** Herbivory, Avian predation, Bird communities, Native oak forest, Connectivity, Afforestation

## Abstract

The value of novel native broadleaf woodlands for biodiversity conservation is important to consider for adequate forest management in rural landscapes. Passive reforestation has been proposed as a cost-efficient tool for creating networks of novel native forest stands that would help restoring biodiversity and associated ecosystem services. Yet to date the ecological functioning of such stands remains strongly understudied compared to forest remnants resulting from longer-term fragmentation. We assessed how the size and connectivity of newly established Pedunculate oak (*Quercus robur* L.) stands in rural landscapes of SW France affect rates of herbivory by different insect guilds as well as rates of avian insectivory and the abundance and richness of insectivorous birds. Comparing 18 novel forest stands along a gradient of size (0.04-1.15 ha) and cover of broadleaf forests in the surroundings (0-30% within a 500 radius), we found that even the smallest stands are colonised by leaf miners and chewers/skeletonizers, and that rates of herbivory are globally comparable to those reported from older and larger oak forests. The size of stands had a relatively minor effect on herbivory, whereas it increased the abundance of insectivorous bird. It also determined rates of avian insectivory as estimated by an experiment with plasticine caterpillars. These rates were however rather low and unrelated with the extent of herbivory in the stand. Overall, our study indicates that insect herbivores tend to react more rapidly to the establishment of novel native forests than their avian predators as the latter may depend on the development of larger patches of suitable habitat in the surrounding landscape. To favour a rapid build-up of diverse, and hence stable, trophic networks involving insect herbivores and their predators, woodland creation schemes should therefore primarily focus on habitat size and quality.

## Introduction

Forest fragmentation is well-known to alter patterns of species distribution and abundance, relationships between organisms and resulting ecosystem processes (Ewers and Didham, 2006; Fahrig, 2017; Haddad et al., 2015; Lindenmayer and Fisher, 2013). Among others, it exerts strong effects on trophic cascades such as plant-herbivore-predator interactions, eventually affecting rates of tree damage and health (Bagchi et al., 2018; Chávez-Pesqueira et al., 2015; Rossetti et al., 2017). While forest fragmentation continues to occur in many regions of the world, forest cover is increasing in many others as a consequence of active planting and passive afforestation following rural abandonment (Fuchs et al., 2015; Hansen et al., 2013). For instance, Europe has experienced a steady increase of forested surfaces by 0.8 million ha per year since 1990 (Forest Europe, 2015), a trend that is expected to continue in the coming decades (Fuchs et al., 2015; Schröter et al., 2005). Habitat defragmentation through passive afforestation has been proposed as an effective tool to reinforce biodiversity and ecosystem functioning in rural and urban landscapes where forest stands were formerly sparse and isolated (Fischer et al., 2006; Rey Benayas et al., 2008; Rey Benayas and Bullock, 2012). Yet little ecological research has to date focused on newly established native forest stands and we largely ignore whether trophic interactions in such stands underlie similar mechanisms as in remnants of similar sizes but resulting from forest fragmentation.

Novel native forest stands establish from a few founder trees that colonize an available habitat patch within an unsuitable matrix through long-distance dispersal and fill their neighbourhood with their offsprings (Gerzabek et al., 2017; Sezen et al., 2005). Such stands share certain characteristics that set them apart from those created by fragmentation: (i) they typically are quite small-sized – even smaller than the smallest fragments of remnant forest; (ii) they are dominated by young trees, resulting in a reduced amount and range of habitats available to forest-dwelling species (Franklin, 1988; Fuller et al., 2018); and (iii) all their species necessarily originate from colonization events over a limited period of time, implying that these systems are triggered by immigration credit instead of extinction debt (Jackson and Sax, 2010). Recent studies on insect and bird species richness along chronosequences of novel native forest development have shown that these are rapidly colonized by woodland generalists whereas specialists can still remain absent even 150 years after forest establishment (Fuentes-Montemayor et al., 2015; Fuller et al., 2018; Whytock et al., 2018). These studies also revealed that local stand characteristics are relatively more important than landscape characteristics for successful colonization by insects and birds. Similar findings have been reported for planted forests (reviewed in Burton et al., 2018). However, their consequences for trophic relationships between plants, insect herbivores and insectivores remain unknown.

Despite the differences between novel native forest stands and remnant forest fragments, the ecological mechanisms underlying trophic cascades involving trees, insect herbivores and birds can to some extent be inferred from fragmentation studies. These have documented that the size and connectivity of forest stands can shape trophic cascades very differently depending on the relative importance of the bottom-up and top-down effects involved (De La Vega et al., 2012; Rossetti et al., 2014). Thus, small and isolated forest stands provide less and possibly lower-quality resources to herbivores (Chávez-Pesqueira et al., 2015) and their colonization requires longer-distance movements that increase energetic and fitness costs (O’Rourke and Petersen, 2017), eventually resulting in lower herbivore abundance (De La Vega et al., 2012; Simonetti et al., 2007). However, small stands also experience greater edge effects which typically go along with increased herbivory (Bagchi et al., 2018; De Carvalho Guimarães et al., 2014). On the other hand, insect herbivores are more likely to colonize small but closer novel forest stands while their predatory vertebrates are more likely to colonize more distant but larger ones (Barbaro et al., 2014; Bereczki et al., 2014; Cooper et al., 2012; Maguire et al., 2015).

There is broad consensus that, generally, predators can notably reduce insect herbivory by regulating herbivore populations (Böhm et al., 2011; Letourneau et al., 2009; Maguire et al., 2015; Rosenheim, 1998). However, their actual relevance in novel native forest stands depends strongly on how both prey and predators respond to stand size and connectivity (Gripenberg and Roslin, 2007). This study investigated how levels of insect herbivory, avian predation and the abundance and diversity of insectivorous birds in recently established native Pedunculate oak (*Quercus robur*) forest stands are influenced by their size and the cover of broadleaf forest in the surrounding landscape. Specifically, we addressed the following questions: (i) Does herbivory increase or decrease along gradients of increasing stand size and connectivity? (ii) Does avian predation increase or decrease along the same gradients? (iii) Are the observed trends related with the local abundance and diversity of insectivorous birds? We contrast our findings with those reported from studies of forest fragmentation and discuss implications in a context of increasing forest connectivity following ongoing changes in landscape use and management (Burton et al., 2018; Rey Benayas and Bullock, 2012).

## Material and methods

### Study area and selection of study sites

The study was carried out in the Landes de Gascogne region (south-western France) about 40 km southwest of Bordeaux (44°41’N, 00°51’W). The region is characterized by an oceanic climate with mean annual temperature of 12.8°C and annual precipitation of 873 mm over the last 20 years. The area is covered by extensive plantations of maritime pine (*Pinus pinaster* Ait.) interspersed with small stands of broadleaved forests that are dominated by Pedunculate oak (*Quercus robur* L.) and contain Pyrenean oak (*Quercus pyrenaica* Willd.), birch (*Betula pendula* L.) and other tree species in minor abundance. Such stands are largely exempt from forest management. Many are actively expanding (Gerzabek et al., 2017), favoured by a recent change in silvicultural management that tends to conserve broadleaved trees recruiting within adjacent pine plantations as a mean of conservation biological control (Castagneyrol et al., 2014; Dulaurent et al., 2011).

We carefully selected a total of 18 novel oak forest stands along gradients of stand size and connectivity (Fig. A1). To ensure that forest stands were of recent origin, we confirmed on aerial photographs from the 1950s that only very few trees were present at that time. We measured the stand area (henceforth referred to as stand size) as the minimum polygon including all oak trees with a stem diameter at breast height of ≥3cm (range: 0.04-1.15 ha; Table A1). The basal area of the stand was also measured and was highly correlated with stand size so we decided to include only stand size in the analysis (Pearson *r* = 0.92, *P* < 0.05). We quantified the spatial connectivity of stands to more ancient forests by calculating the cover of broadleaf forests in a circular buffer of 500 m radius around each stand (range: 0-30%). The size of the buffer (78.5 ha) has previously been shown to be well-suited for studying plant-herbivore-predator interactions (Barbaro et al., 2014; Chaplin-Kramer et al., 2011). Habitat mapping was based on aerial photos using QGIS version 2.18.13 (Quantum GIS Development Team, 2017). Stand size and connectivity were not correlated (Fig. A1; Pearson *r* = 0.39, *P* = 0.11).

### Leaf insect herbivory

In early June 2017, we haphazardly selected four adult oak trees in each forest stand for assessing herbivory and avian predation. On each tree, we cut two south facing and two north facing branches, respectively, at 4 and 8 m height and sampled 20 fully developed leaves from each branch (summing 80 leaves per tree and 320 per stand). Leaves were taken to the laboratory for counting the number of leaf mines and galls per leaf and for estimating the percentage of leaf surface consumed or scratched by chewing and skeletonizing herbivores. A previous study (Giffard et al., 2012) had shown that the most common chewers and skeletonizers in the study area are Lepidoptera and Hymenoptera (sawfly) larvae. We distinguished eight levels of surface damage (0%, 1-5%, 6-10%, 11-15%, 16-25%, 26-50%, 51-75%, and >76%). The gall records were finally discarded from the study because they were too infrequent for independent analyses. In the following, we will refer to ‘herbivory’ as the tree level average leaf area removed by chewing or skeletonizing invertebrates, and to ‘number of mines’ as the average number of mines per leaf. We used the number of mines instead of the proportion of leaves with mines as 9 % of leaves had more than one mine.

### Avian predation

We used plasticine caterpillars made of plasticine (Staedler, Noris Club 8421, green[5]) to estimate predation on insect herbivores. Although not representative of absolute predation rates in the wild, this method allows to compare relative avian predation across stands (González-Gómez et al., 2006; Gunnarsson et al., 2018; Lövei and Ferrante, 2017). Plasticine caterpillars were 30 × 3 mm and light green to mimic late-instar larvae of caterpillars commonly found on oak in the field (Barbaro et al., 2014). We secured 10 plasticine caterpillars at 1.5-2 m height in the canopy of each of our four experimental trees per stand using 0.5 mm metal wires. Predation on plasticine caterpillars was surveyed every six to eight days from 15th May to 15th June (Low et al., 2014). Previous studies have shown that this time period matches the peak activity of insectivorous birds in the study area and is therefore relevant to quantify variation in avian predation (Barbaro et al., 2014; Bereczki et al., 2014; Castagneyrol et al., 2017). All caterpillars with beak marks left by insectivorous birds were recorded and replaced with undamaged ones during each survey. We decided to discard marks putatively left by insectivorous arthropods because we did not assess insectivorous arthropod communities of the stands (see below for birds). Previous to statistical analysis, we standardized our observation by calculating the mean daily predator activity per tree.

### Bird communities

We surveyed the insectivorous bird community in each forest stand using 10-min point counts. Censuses were performed by a trained observer between 6:00 and 9:00 a.m. from the centre of the stand. Each stand was censused twice, once between 26th May and 2nd June and a second time between 21th and 29th June during the exposure period of plasticine caterpillars. All birds within the stand were recorded. Further analysis considered only those species that have a predominantly insectivorous diet during the breeding season. We used the highest count of a given species during any of the censuses as estimate of its abundance within the stand.

### Data analysis

We built three types of models for our different response variables. First, we used linear mixed-effect models (LMM) to model either insect herbivory or the number of mines as a function of stand size (’Size’), stand connectivity in the surrounding landscape (‘Connectivity’) and their interaction (‘Size × Connectivity’). Size, Connectivity and Size × Connectivity were included as fixed effects and the identity of the stand as a random factor. With these predictors three different models were built, each with one further fixed effect, to assess the influence of insectivorous birds on herbivory. These additional fixed effects were either predation on plasticine caterpillars (measured experimentally) or the abundance or species richness of insectivorous birds in the stand (recorded during point counts). We analysed these effects separately because of their non-independence. Second, we modelled predation on plasticine caterpillars as a function of stand size, stand connectivity and their interaction. All were included as fixed effects and stand identity as random effect. Adopting the same approach as for herbivory and the number of mines, we built three models with either herbivory or the abundance or species richness of insectivorous birds per stand as additional fixed effect. Third, we built a generalised linear model (GLM) with stands as replicates to assess the effect of stand size, connectivity and their interaction on the abundance and richness of insectivorous birds. We used Quasi-poisson and Poisson error distributions to model bird abundance and bird species richness, respectively.

All continuous predictor variables were scaled and centred prior to modelling to make their coefficients comparable (Schielzeth, 2010). We first built a full model including all fixed effects, interactions and random factors. Then we applied model simplification by sequentially removing non-significant fixed effects, starting with the least significant interaction. We stopped model simplification with the minimum adequate model when all non-significant terms were taken out. Hereafter, we only report statistics for the simplified models. We estimated and compared model fit by calculating marginal and conditional R^2^ (respectively R_m_^2^ and R_c_^2^) in order to estimate the proportion of variance explained by fixed (R_m_^2^) and fixed plus random factors (R_c_^2^) (Nakagawa and Schielzeth, 2013).

All analyses were done in R version 3.4.1 (2018), using the following packages: *car, doBy, forecast, lmerTest, MuMIn* and *vegan* (Barton, 2018; Fox and Weisberg, 2011; Højsgaard and Halekoh, 2018; Hyndman et al., 2018; Kuznetsova et al., 2017; Oksanen et al., 2018).

## Results

Insect herbivory was on average (± se) 8.02 ± 4.51 % (Table A1). The effect of stand size on herbivory depended on the connectivity of the stand (significant Size × Connectivity interaction, Table 1): herbivory tended to increase with stand size in landscapes with a low stand connectivity whereas it decreased in landscape where broadleaf forests where more abundant (Fig. 1). Neither avian predation on plasticine caterpillars nor bird abundance or richness had a significant effect on herbivory. The number of mines per leaf was on average 0.07 ± 0.05 (Table A1) and decreased in stands that were more connected. Leaf miners were not affected by stand size (Table 1).

**Table 1.** Summary of LMM testing the effect of stand size, connectivity, their interaction and either Avian predation, abundance or richness on insect herbivory. For avian predation the effect of stand size, connectivity, their interaction and either herbivory, bird abundance or richness were tested. Significant variables are indicated in bold. Only predictors retained after model simplification are shown. Predictors were scaled and centred. *R*^*2*^*m* and *R*^*2*^*c* correspond to the variance explained by fixed and fixed plus random factors, respectively.

| Response | Predictors | $\chi^2$ | Df | Coef. ± SE | *P* | $R^2_m$ ($R^2_c$) |
| --- | --- | --- | --- | --- | --- | --- |
| Herbivory | Size | 0.06 | 1 | 2.010 ± 1.061 | 0.807 | 0.20 (0.43) |
| | Connectivity | 0.87 | 1 | -0.041 ± 0.784 | 0.351 | |
| | Size × Connectivity | 8.35 | 1 | -2.933 ± 1.015 | **0.004** | |
| No. of mines | Connectivity | 4.53 | 1 | -0.016 ± 0.007 | **0.033** | 0.10 (0.31) |
| Avian predation | Size | 3.94 | 1 | 0.135 ± 0.068 | **0.047** | 0.06 (0.13) |

**Fig. 1.**
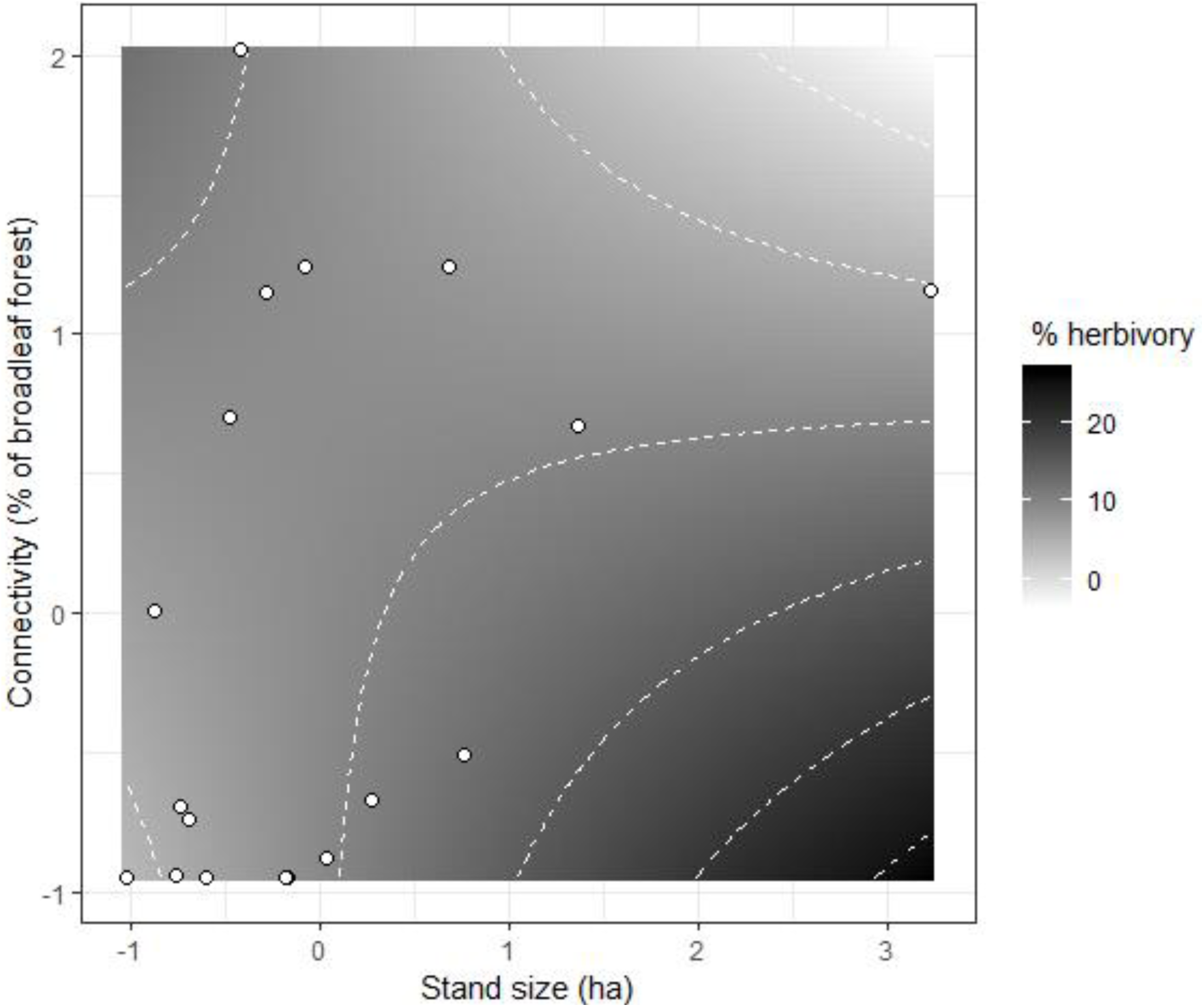
Interactive effect of stand size and connectivity on herbivory. White to black colour scale and isolines show the predicted percentage of herbivory along standardized gradients of stand size (measured as the stand area) and stand connectivity (measured as the cover of broadleaf forest within a buffer of 500 m radius). White dots show the distribution of the original data.

A total of 18 caterpillars out of the 720 exposed (2.5 %) presented marks of bird attacks. Avian predation slightly increased with stand size while it did not vary with stand connectivity or the abundance or richness of insectivorous birds in the stand (Table 1).

We detected a total of 17 bird species within the studied oak stands. The mean (± se) abundance was 4.22 ± 2.59 individuals (range: 1 - 9) and the mean species richness was 3.22 ± 1.66 (range: 1 - 6). The most abundant bird species were blue tit (*Cyanistes caeruleus*), common chaffinch (*Fringilla coelebs*) and chiffchaff (*Phylloscopus collybita*) (Fig. A2). These three species accounted for 38.2 % of all records. Total bird abundance increased with stand size (Fig. 2a, Table 2) and decreased with stand connectivity (Fig. 2b, Table 2). The strength of stand size and connectivity effects was comparable although their effects were opposite. Species richness did not vary with stand size nor with stand connectivity.

**Table 2.**
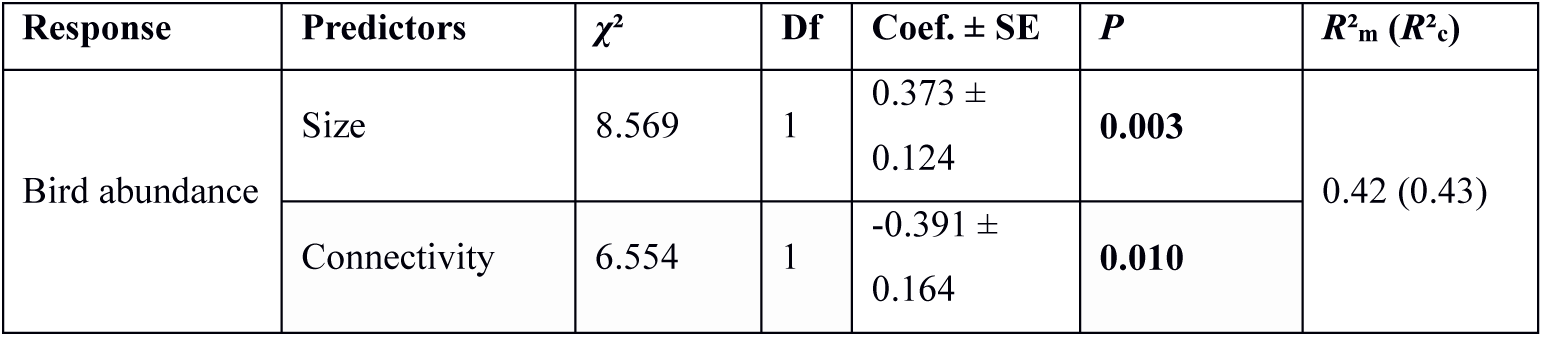
Summary of the GLM on insectivorous bird abundance and species richness as a function of stand size and connectivity. Only predictors remaining after model simplification are shown. Stand size and connectivity were previously standardized. LR: Likelihood Ratio.

| Response | Predictors | $\chi^2$ | Df | Coef. ± SE | *P* | $R^2_m$ ($R^2_c$) |
| --- | --- | --- | --- | --- | --- | --- |
| Bird abundance | Size | 8.569 | 1 | 0.373 ± 0.124 | **0.003** | 0.42 (0.43) |
| | Connectivity | 6.554 | 1 | -0.391 ± 0.164 | **0.010** | |

**Fig. 2.**
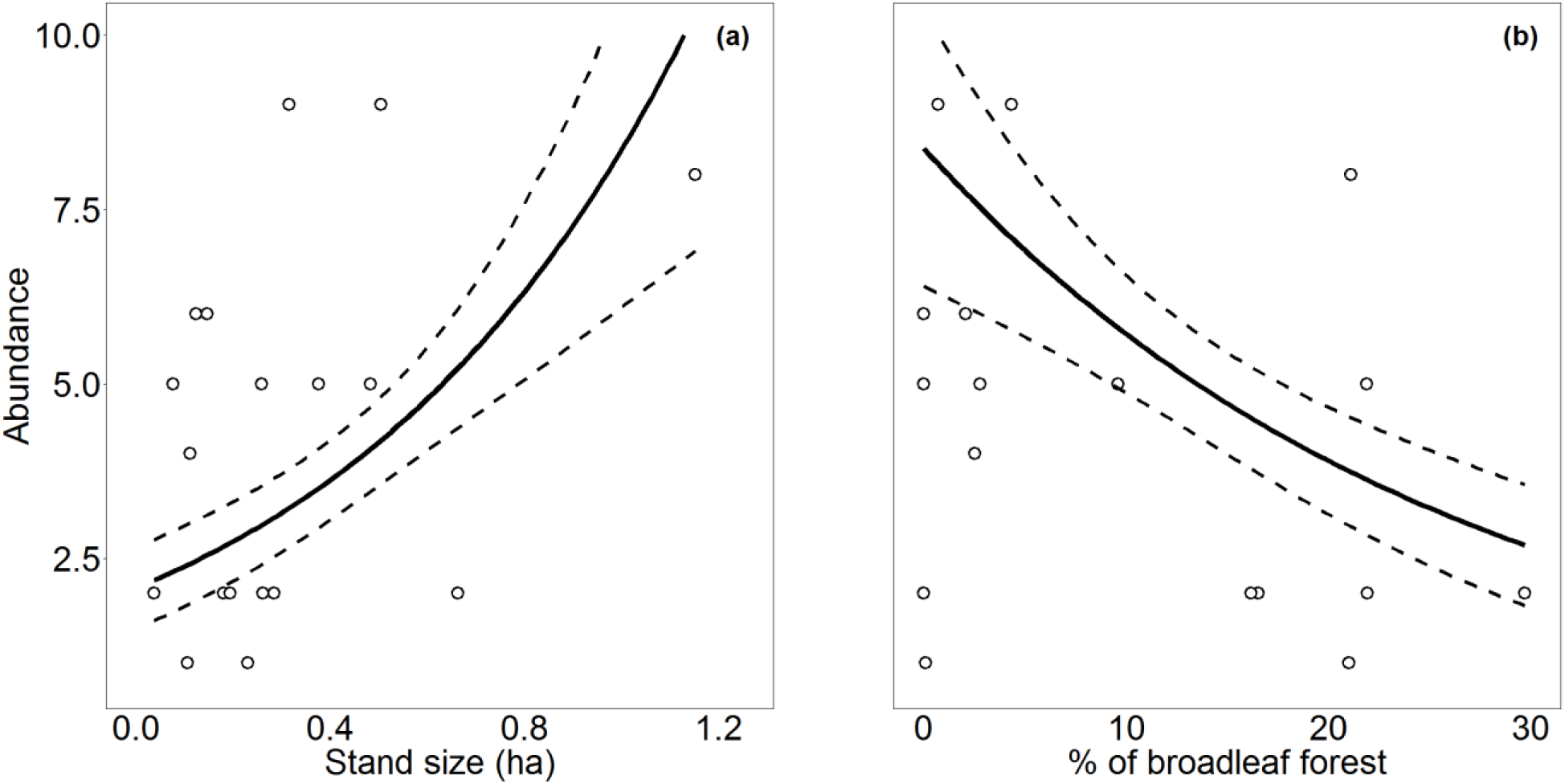
Effects of stand size and connectivity on bird abundance (a, b). Dots represent the individual stands. Solid lines and dashed lines represent model predictions and corresponding standard errors, respectively.

## Discussion

Our study revealed that the size and connectivity of novel native forest stands affect herbivorous insects and insectivorous birds in different ways. While the abundance of leaf miners depended on stand connectivity alone, herbivory by chewers and skeletonizers was influenced by an interplay between stand size and connectivity, and bird abundance (but not species richness) showed consistent independent and opposite responses to stand size and connectivity. This divergence of relationships is likely to arise from differences in the spatial grain of habitat perception and use by the different trophic guilds. It illustrates the complex nature of trophic cascades involving trees, insect herbivores and insectivorous birds in novel native forest stands (Gripenberg and Roslin, 2007).

### Insect herbivores

The observed decrease in the abundance of leaf mining insects with increasing stand connectivity contrasts with previous detailed studies of leaf miners on *Quercus robur* (Gripenberg et al., 2008; Tack et al., 2010) that reported the opposite trend. Importantly, however, these studies focused on a finer spatial grain since they compared individual oak trees with different small-scale ecological neighbourhoods, not with entire forest stands. While the context of their study implies limited movement ranges of leaf mining insects, our results suggest that low abundance of source populations in the surroundings does not limit the ability of this guild to colonise and persist in small novel forest stands. The observed trend could instead be triggered by a resource dilution effect (Otway et al., 2005) whereby herbivore concentrate on the fewer available host individuals (Bañuelos and Kollmann, 2011). Dietary specialists such as many leaf miners should be particularly concerned by resource dilution (Elzinga et al., 2005).

Herbivory by chewing and skeletonizing insects was triggered by stand size in areas where oaks were generally sparse. Positive relationships between stand size and herbivory have also been reported by several studies conducted in considerably larger forest fragments (De La Vega et al., 2012; Simonetti et al., 2007 but see Maguire et al., 2015; Silva and Simonetti, 2009). They could arise from a higher density and/or diversity of insect herbivores in larger stands (Chávez-Pesqueira et al., 2015), as predicted by the resource concentration hypothesis (Hambäck and Englund, 2005; Root, 1973). This hypothesis states that the intensity of physical and chemical cues makes these stands more likely to be found and colonised and less likely to be left by herbivores. The resource concentration hypothesis should be particularly relevant in small habitat patches, such as those of our study system. However, we found that leaf herbivory ceased to increase with stand size and started instead to decline when broadleaf forest became more abundant in the surroundings. We have two possible, non-exclusive explanations for this phenomenon: (i) colonization rates of chewers and skeletonizers could generally be so high in our study system that even the smallest forest stands will be effectively reached (and, if necessary, re-colonized) when a certain threshold abundance of suitable habitats and associated herbivore source populations exist in the landscape (Fahrig, 2013). This hypothesis is supported by the fact that novel established forest stands are very rapidly colonised by woodland generalist species (Fuentes-Montemayor et al., 2015; Fuller et al., 2018). Second, (ii) insect herbivory tends to be favoured by edge effects (De Carvalho Guimarães et al., 2014), especially when it involves generalist species (Bagchi et al., 2018). Edge effects decrease in larger stands, which would counteract other positive effects of stand size on herbivory. Both explanations together suggest that the patterns of leaf herbivory that we observed are likely to be primarily driven by a relatively limited set of mobile generalist species. These species generated however leaf consumption rates that were low but comparable to those recorded in many older and larger oak forests (Gunnarsson et al., 2018; Moreira et al., 2018; Sanz, 2001), and they enabled a quick build-up of trophic cascades even in the smallest and youngest stands of our study system (Hagen et al., 2012).

### Avian insectivores and insectivory

Overall bird abundance and species richness were rather low as well as the size of the stands compared to previous works conducted in the same area (Barbaro et al., 2005; Giffard et al. 2012), and so was also the rate of avian predation (Castagneyrol et al., 2017). Previous studies by Genua et al. (2017), Peter et al. (2015) and Ruiz-Guerra et al. (2012) also found an increase in bird abundance with an increase in continuous forest in the landscape. These forests were however larger than the stands of our study, supporting the idea that avian predation rate and bird abundance (but not species richness) increased with stand size. Overall, these findings suggest that the activity of insectivorous birds in our study system is constrained by the carrying capacity of their wooded habitats. Typical breeding season territories of the most frequently recorded bird species actually exceed the size of our smallest stands (Hinsley et al., 1995) and only the largest stands could regularly sustain more than one territory of the same species. These large stands should also provide the broadest range of tree ages and vegetation structures to different species, although it certainly is still inferior to that of mature forests (Fuentes-Montemayor et al., 2015). Habitat diversity and quality might then also be behind our rather surprising finding that bird abundance (although not species richness) tended to decrease with increasing stand connectivity (Fig. 2). Around the least connected stands, the broadleaf forest cover typically consisted of small, early-successional woodland patches, whereas several of the most connected stands were close to more continuous, older forests, expected to host a large functional diversity of insectivorous birds. The habitat quality of our focal stands should hence equal or exceed that of their surroundings in the former case but be inferior in the latter. The lower use of stands located near larger forests could then be interpreted as a resource dilution effect (see also Berg, 1997; Brotons et al., 2003). That we failed to see this landscape-scale effect reflected in our predation experiment could then simply be due to the low overall number of caterpillar attacks that we recorded and/or other potential limitations of the experimental approach (Muchula et al., 2019). It is however consistent with previous studies that fail to correlate herbivory with predation on plasticine caterpillars (Bereczki et al., 2014; Castagneyrol et al., 2017).

### Tree-herbivore-insectivore interactions and the management of novel native forests

To date most studies on the ecological impacts of active or passive afforestation in fragmented landscapes have focused on patterns of biodiversity (reviewed in Burton et al., 2018), whereas functional ecological aspects have received far less attention (but see Rey Benayas and Bullock, 2012). Our study on bird-insect relationships in novel established native forest stands adds a novel perspective to this field. Taken together, our results indicate that novel forest stands can be very effectively colonised by different guilds of insect herbivores. Although this process is likely to involve primarily a subset of mobile generalist species, these alone can generate levels of herbivory that are quite comparable to those at later stages of forest succession and in areas with higher forest cover. In turn, the build-up of insectivorous bird communities tends to occur more slowly because these depend more than their prey on the development of suitable habitat patches of a certain minimum size (Genua et al., 2017). Birds, as long-lived mobile vertebrate insectivores, typically need to find enough substitutable or non-substitutable resources in the surrounding habitat patches to fulfil entirely their life cycles, namely landscape supplementation and complementation processes (Brotons et al., 2005; Dunning et al., 1992; Fahrig, 2017; Tubelis et al., 2004). Globally, we failed to detect any evidence of top-down control of herbivory by predators. As a consequence, trophic networks in our study system are likely to underlie strong stochasticity, resulting in extensive among-stand heterogeneity and variation through time, which is also typical of forest ecosystems having experienced long-term fragmentation processes (Hagen et al., 2012; Bregman et al., 2015; Fahrig, 2017).

The value of native broadleaf woodlands for biodiversity conservation is important to consider for sustainable forest management in rural landscapes. Landscape defragmentation through networks of novel native forest stands represents a cost-efficient tool for restoring biodiversity and numerous associated ecosystem services (Rey Benayas and Bullock, 2012). Yet the dynamics and ecological functioning of novel native forest stands remain much less well understood than those of forest remnants resulting from fragmentation. Our study underpins that different trophic guilds respond very differently to these novel habitats depending on the spatial grain at which they perceive and exploit them (Gripenberg and Roslin, 2007). To favour a rapid build-up of diverse, and hence stable, trophic networks involving insect herbivores and their predators, woodland creation schemes should focus on habitat size and quality rather than connectivity, including a management that facilitates a diverse tree and understorey vegetation structure (see also Burton et al., 2018; Fuller et al., 2018).

## Acknowledgements

We thank Christophe Poileux, Victor Rébillard, Fabrice Vetillard and Elias Garrouj for their technical assistance in the field and in the laboratory. E.V.C was founded by the project SPONFOREST (grant BIODIVERSA 2015-58).

## Author contributions

E.V.C., B.C., A.H and I.V.H conceived the study and acquired the data. E.V.C and B.C analysed the data. E.V.C., B.C and A.H drafted the first version of the manuscript. All authors wrote the final version of the manuscript.

