## Supplementary material for "Insect herbivory and avian insectivory in novel native oak forests: divergent effects of stand size and connectivity": Suplementari Fig. A1 and A2, table A1

### Appendix A

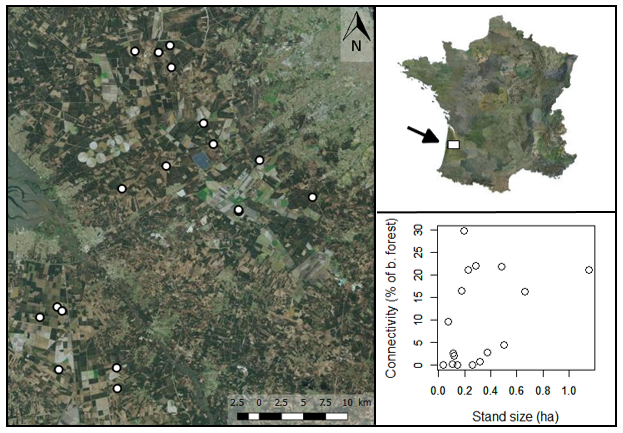

**Fig. A1.** Location map of the study area in the Aquitaine region, south-western France, showing the 18 oak stands at the top right and left of the figure, and figure showing stand size (ha) and connectivity of each stand at the bottom right of the figure.

**Table A1.** Information about the location and size of the oak stands included in the study and summary of the results of herbivory (% leaf damage and Number of mines), predation on plasticine caterpillars and bird abundance and species richness within each stand.

| **Stand** | **Latitude** | **Longitude** | **Stand size (ha)** | **No. of Oaks** | **Herbivory** | **No. of mines / leaf** | **Avian predation on caterpillars** | **Bird abundance** | **Bird species richness** |
| --- | --- | --- | --- | --- | --- | --- | --- | --- | --- |
| 1 | 44.743 | -0.800 | 0.375 | 110 | 13.88 | 0.147 | 0.589 | 5 | 4 |
| 2 | 44.729 | -0.733 | 0.123 | 28 | 4.68 | 0.078 | 0.089 | 6 | 4 |
| 3 | 44.764 | -0.816 | 0.179 | 35 | 13.72 | 0.056 | 0.000 | 2 | 1 |
| 4 | 44.568 | -1.011 | 0.315 | 50 | 4.41 | 0.069 | 0.268 | 9 | 6 |
| 5 | 44.564 | -1.004 | 0.111 | 32 | 6.58 | 0.050 | 0.000 | 4 | 3 |
| 6 | 44.556 | -0.035 | 0.106 | 30 | 6.50 | 0.059 | 0.268 | 1 | 1 |
| 7 | 44.834 | -0.919 | 0.504 | 33 | 13.77 | 0.025 | 0.324 | 9 | 6 |
| 8 | 44.834 | -0.885 | 0.229 | 71 | 5.81 | 0.044 | 0.893 | 1 | 1 |
| 9 | 44.842 | -0.869 | 0.663 | 132 | 6.54 | 0.034 | 0.491 | 2 | 2 |
| 10 | 44.819 | -0.865 | 0.483 | 150 | 4.92 | 0.056 | 0.000 | 5 | 4 |
| 11 | 44.677 | -0.760 | 0.261 | 55 | 7.31 | 0.141 | 0.263 | 2 | 2 |
| 12 | 44.675 | -0.759 | 0.036 | 17 | 5.58 | 0.103 | 0.781 | 2 | 2 |
| 13 | 44.693 | -0.655 | 0.146 | 64 | 3.97 | 0.072 | 0.179 | 6 | 4 |
| 14 | 44.504 | -0.004 | 0.193 | 43 | 10.55 | 0.022 | 0.179 | 2 | 2 |
| 15 | 44.692 | -0.928 | 1.151 | 156 | 6.35 | 0.088 | 0.964 | 8 | 6 |
| 16 | 44.719 | -0.869 | 0.283 | 29 | 11.65 | 0.066 | 0.536 | 2 | 2 |
| 17 | 44.509 | -0.922 | 0.075 | 16 | 8.78 | 0.072 | 0.089 | 5 | 4 |
| 18 | 44.487 | -0.920 | 0.258 | 38 | 9.40 | 0.075 | 0.655 | 5 | 4 |

**Fig. A2.** Proportion of the total abundance of each insectivorous bird species recorded during the study. The total number of individuals per species is indicated on each bar.

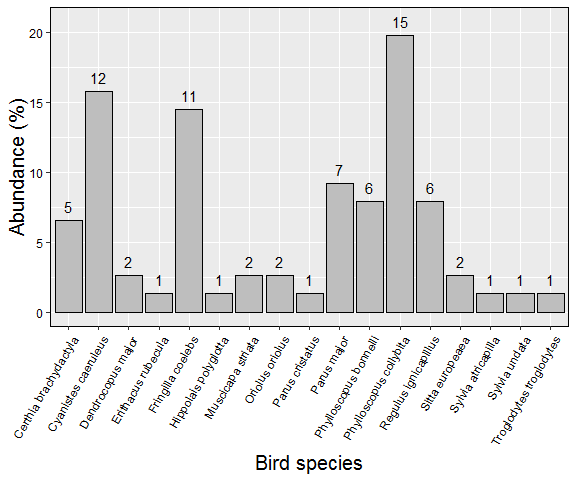
